# Improved BioCyc Operon Prediction: Revisiting the Operon Prediction Problem

**DOI:** 10.1101/2024.06.23.600222

**Authors:** Peter E. Midford, John Cadigan, Peter D. Karp

## Abstract

**Introduction:** Operon prediction is a valuable component of microbial-genome annotation because operon organization can yield inferences about gene function, and because knowledge of operon structure can aid the interpretation of gene expression data.

**Methods:** We present a number of improvements to the existing Pathway Tools operon predictor based mostly on 7 new features that we hypothesized would increase its performance. The new features include shared Gene Ontology biological process terms, similarity of codon usage and GC content, correlated gene expression, and shared protein complex.

**Results:** We evaluated the proposed 7 new features and found that the addition of 6 of them improved the performance of the operon predictor from 79.55% to 83.49%, a decrease in error rate of 19.3%. When gene expression data was not included, the accuracy decreased to 82.547, still an improvement of 14.7%. One of the proposed features as well as a previously used feature had no effect and were removed.

**Discussion:** Although some of the new features had strong predictive value individually, when combined with the other features they did not have a large impact on predictive accuracy, suggesting that they were not independent from the other features.

## Introduction

An operon is a set of genes contiguous on a bacterial chromosome that are transcriptionally regulated as a unit. Operon prediction is a valuable component of microbial-genome annotation because genes within the same operon are often functionally related, thus operon organization can yield functional inferences. Furthermore, knowledge of operon structure can aid the interpretation of gene expression data by illuminating sets of co-regulated genes.

The Pathway Tools (PTools) operon predictor was developed in 2004 [1] and has been applied to tens of thousands of bacterial genomes both by the many users of PTools and for the BioCyc.org website. The goal of our work was to modernize this operon predictor by training and validating it on the expanded set of experimentally determined operons now present in the EcoCyc database for *E. coli* K-12 MG1655 by replacing its use of the outdated MultiFun ontology [2] in favor of Gene Ontology (GO); and by evaluating a number of additional features that we hypothesized might improve its performance. In particular, we sought to integrate gene expression data into operon prediction.

We can formulate the operon-prediction problem in several alternative ways depending on what information we assume is available to the predictor, which determines what features the predictor might employ. The simplest formulation is: given a genome sequence, plus a set of coding regions on that sequence (such as predicted by a gene finder), find groupings of contiguous sets of genes transcribed in the same direction that correspond to operons. Put another way, the goal is to classify every pair of adjacent genes in the genome as within an operon, or at a boundary between operons.

A more expansive formulation makes significant additional information available to the predictor, such as the functional annotation of each gene including GO terms, multimeric complexes to which gene products are assigned, metabolic pathways in which gene products participate, and one or more gene-expression datasets for the organism. This formulation is the one used in this work. Since different genomes will have varying amounts of the preceding information present (e.g., gene expression information is not available for every genome, nor are GO annotations), the operon predictor should be robust when some of this information is not available. This formulation also seeks to classify every pair of adjacent genes as within an operon or at an operon boundary.

These formulations simplify operon prediction in two respects. First, they ignore that operons are often controlled by multiple promoters, some of which are internal to the operon and produce transcripts containing a subset of the operon genes. For example, the *E. coli* operon for *thrS* [3] contains seven promoters (most with experimental evidence) and four terminators. Second, these formulations ignore the fact that bacterial genes sometimes (rarely) overlap one another for most or all of their lengths on either the same or opposite DNA strands. For example, the *E. coli* genes *sgrT* and *sgrS* completely overlap one another, and are transcribed in the same direction. How such overlaps affect operon organization is unknown.

## Methods

This operon predictor is based on an existing predictor [1] used in the Pathway Tools software since 2004. Both versions of the operon predictor obtain most input information (e.g., the organism’s genome, gene list, and proteome) from the Pathway/Genome Database (PGDB) describing the organism. The PGDBs are derived from RefSeq entries by the PathoLogic component of PTools [4]. The outputs of the operon predictor are written to the PGDB — each predicted operon is a PGDB object that specifies its component genes and includes a computational evidence code.

The first step performed by both operon predictors is to divide each replicon (chromosome, plasmid, or contig) in the organism’s PGDB into directons — runs of adjacent genes transcribed in one directon. Directons are trivial to compute and are a first step toward computing operons: some directons will themselves be operons (e.g., single-gene directons), whereas other directons will be divided into multiple operons. The code for directon generation is relatively unchanged except that the predictor now supports selecting a subset of the organism’s replicons for operon prediction (the unselected replicons are left unchanged).

### Features for Operon Prediction

In common with other directon-based methods, the PTools operon predictor considers each adjacent or overlapping pair of genes in a directon, and predicts whether the genes in that pair are in an operon or represent a boundary between operons. All features are combined in a logistic regression to classify the pair as either “within an operon” or representing an operon boundary. The set of previous and new features are listed in Table 1.

**Table 1:** Features used by operon predictor. The second column indicates whether the feature was added since the 2004 version of the predictor.

| Feature | New? |
| --- | --- |
| Distance | no |
| Gene Expression | yes |
| Shared GO terms | yes<sup>1</sup> |
| Gene Symbol Similarity | yes |
| Similar Products | yes |
| Same Complex | no |
| Shared Pathway | no |
| Shared Transport | no |
| Up/Downstream shared pathway | no |
| Up/Downstream shared transport | no |
| GC Content | yes |
| Similar Length | yes |
| Codon Usage | yes |

Our training data for developing the new operon predictor consisted of adjacent *E. coli* gene pairs with experimental evidence that either supported or refuted that they were members of an operon. We used this data from EcoCyc version 27.0. If a gene pair was included in an experimentally supported operon (meaning the operon is annotated with an experimental evidence code), we considered that pair to be a positive example. A pair is considered to be a negative example if the pair is within the same operon when one gene in the pair is assigned to an operon with experimental support, but the other gene is not assigned to that operon. Note the evidence codes for the genes themselves are not considered since these codes refer to the functions of the genes and not to their operon assignments.

The features used and/or evaluated for the new operon predictor are as follows.

### Distance

Distance between members of a gene pair was introduced by [5] and is by far the most common feature used in operon predictors [6, 7] as it seems to be one of the strongest predictors of operon membership. Distance refers to the number of nucleotides from the end of one gene’s coding region to the start of the adjacent gene’s coding region. Like our previous predictor, the new predictor estimates the function relating distance and likelihood of operon membership based on *E. coli* data. We have updated the likelihood function based on 18 years of additional curation of known operons into EcoCyc. We took all gene pairs that could be assigned either positive or negative evidence of operon membership and, as in the previous predictor [1], assigned them to bins of width of 10 nucleotides based on the distance between the genes. We computed likelihood ratios based on the counts of positive and negative example gene pairs in each bin (see Figure 1). The previous predictor used a threshold to convert likelihood values into binary features. Since a regression model is used in this predictor, the likelihood ratio corresponding to the gene pair’s distance was entered directly into the regression.

**Figure 1:**
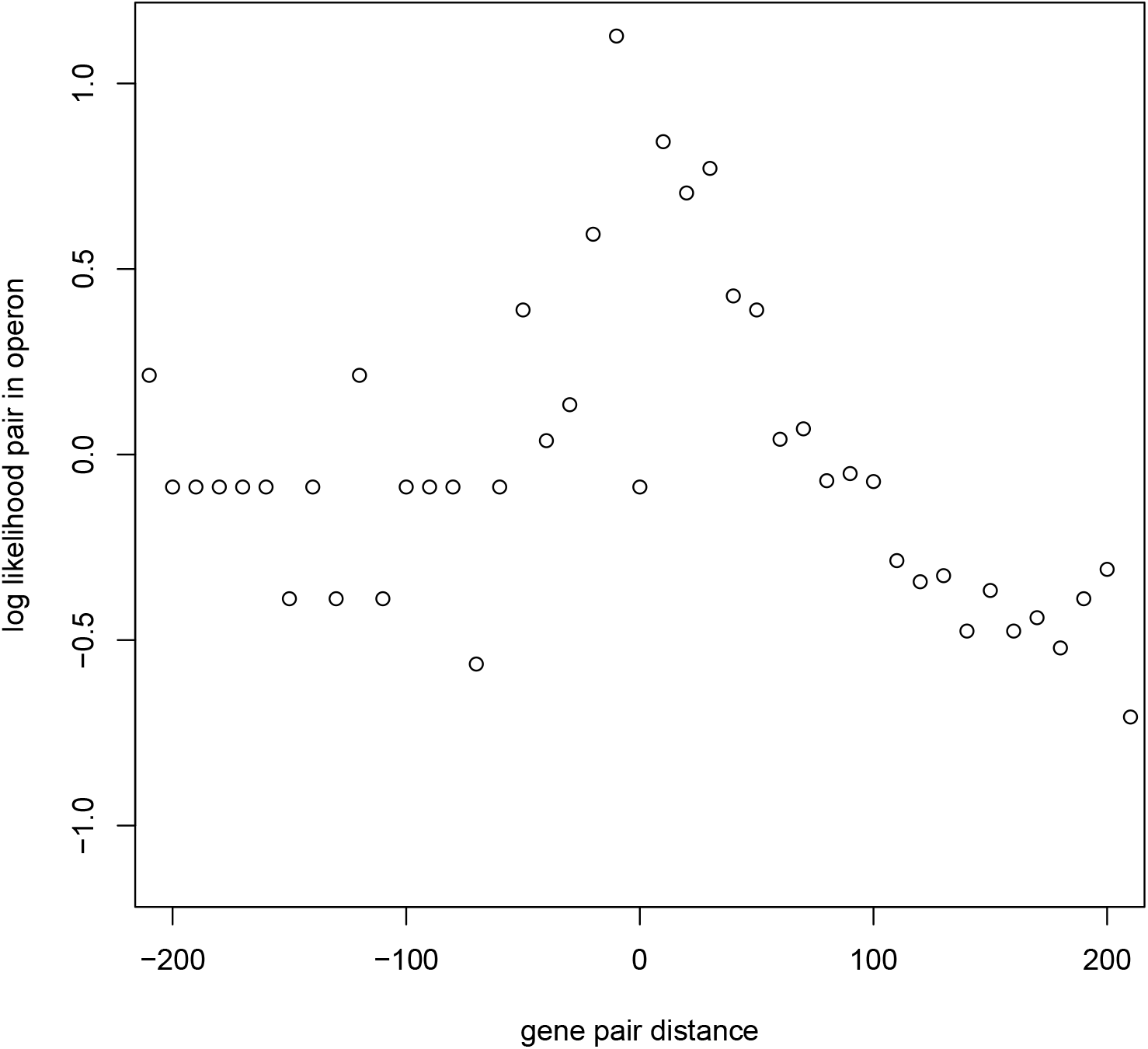
Plot of likelihood ratios of within-operon to boundary pairs versus gene-pair distance; data from Ecocyc 26.5

### Correlated Gene Expression

If gene expression data covering a sufficient number of experimental conditions (*>*= 5) is available, then we use gene expression data as an additional feature. If the Pearson correlation of the gene expression levels of the two adjacent genes is significantly different from zero, it is considered evidence that the pair of genes is in an operon. This is a straightforward way to make use of RNASeq short-read data in which the reads are too short to cover entire transcripts. This feature is used in several operon predictors that make use of transcriptomes [8, 9, 10]. We use a cut-off value of 0.5 for the correlation value — anything less is entered into the regression as 0, values above 0.5 were entered unchanged. If either member of the gene pair has missing data for a condition, we ignore that experiment for the correlation calculation.

To enable use of our operon predictor for organisms for which gene expression data is and is not available, the operon predictor includes one model that includes expression correlation data and a second model that does not. Our user interface allows loading a file of gene expression data that will be used for calculating correlations across multiple experimental conditions.

### Gene Functional Features

If a pair of adjacent genes, A and B, have products P(A) and P(B) with similar biological function, this similarity suggests the genes may be transcribed as an operon. Although several features related to functional similarity were used in our earlier operon predictor, we introduce several new such features here. We also removed the use of the MultiFun ontology to predict functional relatedness because MultiFun has been superseded by GO.

### Gene Ontology Biological Process Overlap

We determine whether gene products have similar function by looking for an overlap in the sets of GO [11] biological process terms for those genes. An overlap is considered evidence that those genes participate in the same process and therefore are likely to be in an operon. The DOOR database [12] uses semantic similarity of assigned GO terms (it isn’t stated which GO sub-ontology was used).

### Protein Complexes

If the products of a pair of adjacent genes are members of a known protein complex, this is another indication the genes are functionally related, thus providing evidence of membership in an operon. We use two types of evidence of membership in a protein complex. The first uses existing assignments of products to complexes in the PGDB, the second uses a lexical comparison of the names of the products of the gene pair. The lexical analysis looks for pairs of product names that match except for substrings such as “subunit A” and “subunit B”. A positive example is the pair of names “hydrogenase 2 large subunit” and “hydrogenase 2 membrane subunit”, whereas the pair “citrate lyase beta subunit” and “citrate lyase acyl carrier protein” would be rejected because the substring “acyl carrier protein” is not a subunit phrase. We did not find other predictors that specifically used membership in complexes as a feature. The earlier version of the predictor [1] implemented this feature using only assignments of products to complexes, but not the lexical analysis of names. We are not aware of other predictors that use this feature.

### Shared Pathway

If the genes in the pair participate in a common pathway, then we consider this form of functional relatedness as evidence the two genes are members of an operon. Numerous other predictors use pathways as features [7, 6, 12], most commonly KEGG pathways. The common pathway feature was used in our previous predictor [1].

### Pathway/Transporter

This rule considers transporters that either import pathway substrates or export pathway products. If one of the genes in the pair is a transporter that either imports or exports a substrate in a pathway for which the other gene contributes an enzyme to, the genes are functionally related, and hence are likely to belong to a common operon. Although this strategy was used in our earlier predictor [1], no other predictor explicitly used a transporter related to pathway feature.

### Similar Product Types

This feature asks whether the products of a pair of adjacent genes are either both proteins or both RNAs. If the products of genes *A* and *B* are both proteins or both RNAs, then we consider that as evidence they are within an operon. We tested a variant of this rule that asks whether the products are both proteins or both regulatory RNAs, but that variant was less effective. We did not find another predictor that used this particular feature.

### Gene Symbol Similarity

This rule considers the gene symbols (short names) assigned to the genes, such as *trpA* and *ileS*. The biological conventions behind assignment of gene symbols imply that genes involved in the same biological system (and thus are functionally related) are usually assigned the same first three letters, with the fourth letter indicating a subsystem or pathway. For example, *E. coli* genes whose names begin with “trp” are all involved in tryptophan metabolism in some way. *trpA–E* are all involved in the biosynthesis of tryptophan, whereas *trpS* codes for tryptophan–tRNA ligase and *trpR* codes for a repressor of the *trpA–E* genes.

The rule we employ is: If the gene symbols for genes *A* and *B* are four characters in length, and the first three characters of the name match, and the first character of the name isn’t “y” and the fourth characters of each name are within a specified lexical distance, then the genes are more likely to be in the same operon.

The exclusion of names with the first character “y” follows a naming convention for *E. coli* genes that assigns such names to genes of unknown function. We experimented with lexical distances of one to six and found that two was the distance that gave the best results. So “trpA” and “trpI” are not considered similar, but “trpA” and “trpC” are.

We did not find other predictors that used this feature.

### Up/Downstream Shared Pathway

When classifying an adjacent pair of genes (e.g., *A* and *B*), if there is evidence other genes in the same directon have functional similarity to either *A* or *B*, it is reasonable to infer that the genes between them are also in that operon. Formally, if a directon contains genes *A, B*, and *C* in that order, and if the predictor is predicting the operon status of the adjacent pair *A* and *B*, and there is evidence that *A* and *C* are in an operon, then it is reasonable to infer that *B* is in that operon. Likewise, if the predictor is considering the adjacent pair *B* and *C*, and if there is evidence that *A* and *C* are in an operon, then it is also reasonable to infer that *B* is also in that operon.

The predictor addresses this possibility by looking at all genes in the directon that are downstream of *B*. Each gene is paired with *A* and evaluated with the test for shared pathways. If the shared pathways test is true for any such pair, it is evidence that *A* and *B* are within an operon. Likewise, each gene in the directon that is upstream of *A* is paired with *B* and evaluated with the test for shared pathways. If the shared pathways test is true for any such pair, it is evidence that *A* and *B* are within an operon. Apart from [1], we did not find any other predictors that applied this feature across genes in a directon.

#### Up/Downstream Pathway/Transporter

This feature performs testing analogous to the previous feature, but uses the functional relationship tested by the Pathway/Transporter feature. Apart from [1], we did not find any other predictors that applied this feature across genes in a directon.

#### Similar Length

If two adjacent genes greatly vary in length, this feature suggests they may not be in the same operon, under the intuition that genes with very different lengths are more likely to not be functionally related. This feature is true if the ratio of the lengths of the genes in the pair is within a factor of four. DOOR [12] and several other predictors [6] also use the ratio between the lengths of the genes in a pair.

#### GC Content

If two adjacent genes have similar GC content, this rule suggests they might be in the same operon. The intuition is that, given two adjacent genes, if one arose through horizontal transfer and one did not, they are unlikely to be functionally related; this situation is more likely to apply to genes with dissimilar GC content. Because the GC content is a continuous value, the values could be used directly in the model, but we chose a to convert this to a binary variable by setting a threshold. We maximized Youden’s index [13] (essentially the sum of sensitivity and selectivity scores) by doing a simple binary search over the 0 -1 range of thresholds. For each threshold value, we calculated the Youden index for the set of 1633 gene pairs with experimental evidence as predicted strictly by the GC content, stopping the search when the difference of Youden scores was less than a tolerance of 1e-6. The value that maximized the score was selected as the threshold. We did not find other predictors using this feature.

#### Codon Usage

Similar to GC content, we reasoned that, given two adjacent genes, if one arose through horizontal transfer and one did not, they are unlikely to be functionally related; this situation is more likely to apply to genes with dissimilar codon usage. For adjacent pairs of protein-coding genes, we considered codon usage as determined by the cosine similarity of the 64 element codon usage vector for the pair of genes. Since this is a continuous measure, we converted it to a binary feature in the same manner as for GC content. The undefined similarity when one or both genes were RNA coding was coded as zero. As with GC content, we made this a binary feature by searching for a value that maximized the Youden index. Fortino et al.[9] used a different codon usage statistic.

### Generating Gene Expression Data

We used previously submitted RNASeq data to generate a set of gene expression data to use in testing the effectiveness of correlated gene expression as a feature. We used data previously used with Rockhopper [8, 10] as we were already using Rockhopper as a reference for an operon predictor that used RNASeq data.

We downloaded 51 RNAseq datasets from the Sequence Read Archive at the National Center for Biotechnology Information relevant to *E. coli* listed in the supplemental material of [8], assigned them to 21 experimental conditions (see SupFile5-Expression-Experiments.txt) based on both growth conditions and genetic background, and used Rockhopper to generate a file of gene expression levels across the 21 experimental conditions (See SupFile4-RH-script, SupFile6-Expression-Experiments.pdf). We pruned extraneous columns from the Rockhopper output to generate a file of gene expression levels across each experimental condition (SupFile3-Ecoli-Gene-Expression.txt). This file of gene expression levels served as the input to our operon predictor.

### Evaluation of Feature Sets

We created datasets consisting of 1633 adjacent gene pairs from EcoCyc that we could assign as either being within the same operon or not based on experimental evidence (see SupFile4-lisp-code.lisp function generate-logistic-data). The datasets (see SupFile1-Pair-Eata-With-Gene-Expression.txt, SupFile2-Pair-Data-No-Gene-Expression.txt) included, for each gene pair with evidence, the experimentally determined within-operon status as well as the values of each feature under consideration. We generated a logistic regression model using R 4.02 [14] and evaluated area under the ROC Curve (AUC) using the ROCR [15] package. The model parameters were then incorporated into a set of weights used by the pair classifier. We evaluated primarily by the model’s accuracy, although we report sensitivity and specificity as well. The AUC statistics were evaluated by subsampling 100 random subsets of 200 data points.

We ran regressions on the full dataset to tune the size of the character window for the Gene Symbol Similarity feature. We considered lexical distances from one to six. We used a simpler method to tune the thresholds for the GC Content and Codon Usage features: these thresholds were assigned by considering those features in isolation and optimizing for the sum of specificity and sensitivity when tested against the 1633 gene pairs with experimental evidence.

## Comparing Other Machine Learning Methods

We used scikit-learn [16] to evaluate several different ML classifiers on the data set that included gene expression (SupFile1-Pair-Data-With-Gene-Expression.txt). We also considered a Conditional Random Field model.

We ran 10-fold stratified cross-validation experiments with other classic machine learning approaches. Logistic regression with the various regularization penalties: L1, L2 and ElasticNet with a L1 ratio of 0.5 [17]. We also tried Support Vector Machines (SVMs) with various kernels: linear, x^2^, and a Radial Basis Function [18]. We also tried a random forest of 10 trees with a maximum depth of 2 [19]. Other hyperparameters were maintained at their default. The set contained 1624 rows of data in which we treated GC Content and Codon Usage as continuous variables (not thresholded), treated the two methods of detecting protein complexes (PGDB data and lexical comparision) as separate features, and included a feature (up/down stream complex) that we later rejected as uninformative. See Table 2. Detailed results from the Conditional Random Field (CRF) model are in the supplementary file SupFile10-CRF.txt

**Table 2:** Various ML methods applied to the problem in 10-fold stratified cross-validation, mean +/- std.

| Model | Accuracy |
| --- | --- |
| l1-logistic-regression | 0.8350 +/- 0.026 |
| l1-logistic-regression-balanced | 0.8220 +/- 0.025 |
| l2-logistic-regression | 0.8350 +/- 0.026 |
| l2-logistic-regression-boosted-5 | 0.8350 +/- 0.019 |
| logistic-regression | 0.8350 +/- 0.026 |
| logistic-regression-elasticnet | 0.8350 +/- 0.025 |
| random-forest | 0.8200 +/- 0.029 |
| svm-linear | 0.8310 +/- 0.023 |
| svm-poly-2 | 0.8270 +/- 0.025 |
| svm-rbf | 0.8200 +/- 0.023 |
| CRF | 0.837 +/- 0.024 |

### Conditional Random Fields

CRFs are discriminatively trained sequence labeling models [20]. Let our sequence of genes X be *X* = *x*_1_, *x*_2_, …, *x*_*n*_ and the tags be *Y* = *y*_1_, *y*_2_, …, *y*_*n*_.. There are *t* possible tags. The CRF model predicts the probability of the sequence of tags given the input: *P* (*Y* |*X*) with unary and transition scores. Unary scores are calculated for each *x*_*n*_ by a linear model for each tag with *U* (*x*_*n*_, *y*_*n*_). The CRF transition scores *T* are a matrix *R*^*txt*^ in which index i,j represents the transition from tag i to tag j. Combining the two together along with the normalization term *Z*(*X*) used to calculate proper probabilities we have:

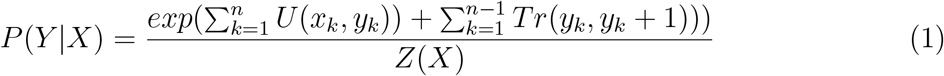

The naive calculation of Z(X) would involve summing up the probabilities over all tag sets, resulting in an *O*(*T*^*N*^) calculation but through the use of the forward algorithm, this is reduced to *O*(*NT* ^2^) through the use of caching intermediate values as a dynamic programming algorithm. The model is trained through minimizing negative log likelihood.

In our sequence labeling Conditional Random Field model [20], we classify individual genes as BEGINNING, INTERIOR, END, or SINGLETON of an operon based on the same features with respect to a preceding gene or a following gene in a given sequence — identical to our classifiers. We experimented with higher order features that included features with respect to more distant genes, but we did not immediately find an improvement. Details of CRF’s are described in section In this paper we use the CRFSuite library with the L-BFGS algorithm to optimize model parameters. We disallow unseen transitions and use default L2 regularization of 1.0.

## Results

### Choice of Machine-Learning Method

We initially selected logistic regression for ease of implementation and updating. After establishing that a logistic regression model would perform at least as well as the existing predictor, we took the *E. coli* experimental operon dataset, and compared logistic regression with a range of other machine learning (ML) methods using scikit-learn [16]. The results, which were consistent with the results of the model fitting, are listed in Table 2

None of the other methods were substantially better than straight logistic regression and several, including variants of SVMs and Random Forest (RF) and several variants of logistic regression, performed worse based on accuracy and AUC scores. At least one other operon predictor [9] considered several machine learning methods, RF and SVMs, but did not include logistic regression.

### Regression

We fit one logistic regression model for *E. coli* that included gene expression data (Table 3) and a second model that did not (Supplementary File SupFile8-NoExpression-Regression).

**Table 3:** Logistic regression coefficients for features from EcoCyc 27.0 and gene expression correlations calculated from 21 RNASeq experiments (Gene Expression). The regression intercept is listed after the features. The mean accuracy across 100 subsamples was 83.0%. The mean AUC ROC was 90.6%

| Feature | Coefficient | p-value | Significance |
| --- | --- | --- | --- |
| Distance | 2.1176 | $2 \times 10^{-16}$ | *** |
| Gene Expression | 1.1329 | $8.88 \times 10^{-8}$ | *** |
| Shared GO terms | 0.5917 | 0.003 23 | ** |
| Gene Symbol Similarity | 1.4070 | $9.89 \times 10^{-7}$ | *** |
| Similar Products | 1.3455 | $4.37 \times 10^{-6}$ | *** |
| Same Complex | 1.4759 | 0.005 28 | ** |
| Shared Pathway | 0.9945 | 0.011 70 | * |
| Shared Transport | 0.7875 | 0.245 50 | NS |
| Up/Downstream shared pathway | 0.3388 | 0.358 24 | NS |
| Up/Downstream shared transport | 0.599 | 0.374 36 | NS |
| GC content | -0.1396 | 0.379 74 | NS |
| Similar Length | -0.4745 | 0.047 05 | * |
| Codon usage | -0.3455 | 0.093 93 | NS |
| Regression Intercept | -1.6155 | $2.26 \times 10^{-8}$ | *** |

### Performance of Features

Ideally, removing all the features R reported as insignificant individually would not have impacted predictor performance. However, removing those features in the *E. coli* model with gene expression resulted in reduced accuracy from 83% to 81% largely due to an increase in false positives. Furthermore, three features (similar length, GC content, and codon usage similarity) were fitted with negative coefficients, suggesting they were providing evidence against operon membership.

To investigate whether any features were actually not predictive of operon membership, we ran simplified regression models consisting of just the feature value and the response. All single feature regressions showed accuracy above 50% and AUC scores above 0.5, however, for the up/downstream transport/pathway feature, the AUC score was only 0.506, and the regression p-value for that feature was not significant. In particular, all three features that had negative regression coefficients in the full model had accuracies between 56% and 64% and AUC scores above 0.53. The best singleton features were gene pair distance (accuracy 78.5% / AUC 0.863) and gene expression correlation (accuracy 73.3 % / AUC 0.789).

### Features We Didn’t Consider

We did not consider expression based RNASeq features, such as expression level continuity or variance, used by [21]. DOOR [12] also used several other gene features: conservation levels of the pair across other genomes, and the presence of certain DNA motifs. Comparative methods, such as conserved gene pairs, clusters of orthologous genes, and phylogenetic profiles (patterns of gene presence across genomes) were also used in other predictors [6].

In addition to the switch from MultiFun to GO function annotations, we removed one existing feature and updated another:

- A feature called upstream/downstream shared protein complex was statistically insignificant and added nothing to the predictive performance of the model.
- The shared protein complex feature added a lexical analysis of product names and also benefited from updates to the protein complex predictor elsewhere in Pathway Tools.

### Performance of the Operon Predictor

Supplementary File SupFile9-Prediction-Stats.txt reports the performance of our predictor for several BioCyc organisms with sufficient numbers of experimentally supported operons. The first row shows results of the full model, which includes the gene expression data for *E. coli*. The second row shows results for *E. coli* using the model fitted without gene expression data. The remaining rows list results for five other organisms that have more than 15 experimentally supported operons. Although the new predictor trends slightly better across these organisms, the small sample sizes make the change in performance difficult to interpret.

As expected, the predictor’s performance agrees fairly well with the performance of the regression model on the full set of gene pairs with experimental evidence. The performance is slightly worse because all pairs with experimental evidence are used in building/evaluating the regression model, but it turns out that not all experimentally supported pairs are considered when pairs are split from directons, so the full predictor’s performance is based on a somewhat smaller set of pairs.

The performance of the predictor on *E. coli* without expression data increased from 79.55% in the old predictor to 82.547% in the new predictor — a decrease in error rate of 14.7%. With expression data the accuracy of the new predictor increased to 83.49% — a decrease in error rate of 19.3%.

### Performance of Other Predictors

We tested the Rockhopper operon predictor (version 2.03 https://cs.wellesley.edu/~btjaden/Rockhopper/) using the same RNASeq data we used for generating our gene expression correlations and the genome files provided by Rockhopper against the set of gene pairs in EcoCyc 27.0 with experimental evidence either for or against being within an operon. The accuracy of Rockhopper’s predictor was slightly better than our previous predictor, though it tended to overpredict within operon pairs (accuracy 80.3%, sensitivity 89.5%, and specificity 68.2%). As distributed, Rockhopper uses only gene expression correlation and distance between pairs to make its pair predictions. Chuang et al. [6] lists a number of operon predictors, and their performance, but differing evaluation data makes it hard to conclude much more than that our results were broadly comparable across accuracy, sensitivity, and selectivity.

## Discussion

The new operon predictor improves both performance and maintainability. The performance improvements are discussed in the results section.

The maintainability improvements will enable us to more easily update the predictor in the future. The previous operon predictor used a complex combination of rules to make predictions. This has been replaced by a standard ML method, and there is a documented procedure for updating the regression model. This is important because the model is derived from EcoCyc data, which is actively curated and will continue to receive updates in the future. In particular, the set of gene pairs with experimental evidence of operon membership will continue to grow, as will other annotations that are relevant to the set of features used by the predictor. There are also procedures for updating the gene expression dataset and the likelihood curve for the gene pair distance data.

### Which Features Mattered?

One surprising result was how many of the features we tried turned out to contribute to model accuracy. This was particularly surprising considering that several features had negative regression coefficients in the model, despite increasing predictive accuracy when considered by themselves. Removal of these features did decrease model accuracy. One explanation for this is that the features, although conceptually different, are not completely statistically independent. Although all features contributed to overall model performance, only some were reported as significant in the regression (Table 3).

Also surprising is that despite the addition of eight new features to the predictor, three of which had very significant p-values individually, the overall accuracy of the predictor was not strongly affected.

The features that mattered most were distance, similar products, gene expression, and similarity of gene symbols. Shared product GO terms, shared product protein complexes, and shared product pathways were significant, but less important contributors.

## Comparing Other Machine Learning Methods

We initially selected logistic regression for ease of implementation and updating. After establishing that a logistic regression model would perform at least as well as the existing predictor and validating features, we took a dataset as described above, and compared logistic regression with a range of other machine learning (ML) methods using scikit-learn [16] and CRF Suite [22]. We found these alternatives have roughly the same level of performance and none stood out relative to a simple logistic regression.

## Supporting information

Supplemental Files ReadMe

Pair Features including Gene Expression

Pair Features without Gene Expression

Gene Expression data

Running the Operon Predictor

Regression weights fit to E coli data without gene expression

Performance of predictor for E coli and five other organisms

Summary of Experiments with Conditional Random Field models

List of NCBI SRR files used for gene expression analysis

Script File for running Rockhopper for correlated expression analysis

R script used to fit the regression model

## Author Contributions

P.E.M wrote the code and performed the data analysis. J.C. performed the comparison of machine learning methods. P.D.K and P.E.M conceived the experiments. P.D.K, P.E.M., and J.C. wrote and reviewed the manuscript.

## Acknowledgments

We thank Robert Landick and Michael Wolfe for valuable discussions.

