## Supplemental Files ReadMe for "Improved BioCyc Operon Prediction: Revisiting the Operon Prediction Problem"

Supplementary Files and How They Were Generated

:::::::::::::::::::::::::::::  SupFile 1  ::::::::::::::::::::::::::

The file SupFile1-pair-data-with-gene-expression.txt contains data for
each gene pair that has experimental information in EcoCyc version 27.0.
This file includes gene expression correlation values as calculated from
RockHopper and code included in our lisp source (SupFile4-lisp-code.lisp).


:::::::::::::::::::::::::::::  SupFile 2  ::::::::::::::::::::::::::

The file SupFile2-pair-data-no-gene-expression.txt contains data for
each gene pair that has experimental information in EcoCyc version 27.0.
This file does not include the gene expression data column; otherwise
this file is the same as SupFile1.

::::::::::::::::::::::::::::::  SupFile3  ::::::::::::::::::::::::::

The file SupFile3-ecoli-gene-expression.txt contains gene expression
data used in our analysis as generated by Rockhopper.
It is based on a set of 51 short
read files from the NCBI Short Read Archive, originally reported in
McClure (2013), specifically those in the E. coli data mentioned
there.  We downloaded the files in .fastq format and generated gene
expression data using the command-line version of Rockhopper 2.03.  We
used the genetics files as provided from Rockhopper's repository of
genome files (http://cs.wellesley.edu/~btjaden/genomes/).  We ran
RockHopper with the script in SupFile5-RH-script.sh. This script
includes the set of SRA files and their grouping into experiments.
The resulting gene expression file contained columns with
intermediate, non-normalized expression levels which we trimmed out to
produce our final file (SupFile3-ecoli-gene-expression.txt).  The
details of the assignment of SRA files to experiments are covered in
SupFile6-Expression-Experiments.pdf.

::::::::::::::::::::::::::::::  SupFile4-RH-Script.sh  ::::::::::::::::::::::::::

The file SupFile4-RH-script.sh contains a script to run RockHopper to
generate the expression data.

::::::::::::::::::::::::::::::  SupFile5-Expression-Experiments.txt  ::::::::::::::::::::::::::

The file SupFile5-Expression-Experiments.txt lists the SRR
expression files downloaded from NCBI SRA using the SRA utilities
that were used to generate the expression data (used in SupFile4-RH-script.sh).


::::::::::::::::::::::::::::::  SupFile6-How-To-Run.txt  :::::::::::::::::::::::::

This file contains instructions for how to run the operon predictor from lisp
in an evaluation mode.  For normal use when building PDGBs see the Pathway Tools
user's guide, included in the Pathway Tools distribution.

::::::::::::::::::::::::::::::  SupFile7-R-Script.R  :::::::::::::::::::::::::

SupFile7 is the R-script used to fit the regression models.  The
important steps are reading the model with read.csv, fitting the model
with glm and retrieving the fit with the summary command. The result
of the summary is copied, pasted and formatted into an assignment to
the appropriate parameter (+logistic-weights+ or
+no-expression-logistic-weights+) in the lisp code.  Note that each
model parameter is a dotted-pair of a parameter name and its
value. Note that each model has a mix of binary and continuous variables
and the binary factor names in the generated model will be suffixed by
'YES'.  Remove those suffixes when updating the parameters.

::::::::::::::::::::::  SupFile8-NoExpression-Regression.txt ::::::::::::::::

This file lists the regression weights for the logistic regression
model generated using EcoCyc 27.0 experimentally supported pair
assignments without gene expression data (c.f. Table 3 in the main paper).

::::::::::::::::::::::::::::::  SupFile9-Prediction-Stats.txt :::::::::::::::::::::::::

This file lists the prediction accuracy, sensitivity and specificity
statistics for E. coli and five other tier 2 organisms. 

::::::::::::::::::::::::::::::  SupFile10-CRF.txt  :::::::::::::::::::::::::

This file summarizes our exploration of Conditional Random
Field models.
