## Supplementary material for "Improved BioCyc Operon Prediction: Revisiting the Operon Prediction Problem": Running the Operon Predictor

 Instructions for replicating reported results

The function evaluate-predict-operons was used to test and generate the results listed in the 
publication.  To run that function:

1. Start the Pathway Tools software that is freely available from SRI for academic  researchers
    and employees of non-profit institutions at https://biocyc.org/download-bundle.shtml.


 2. Once you see a prompt (looks like EC(2):), load the PGDB for the organism of interest.  Use the
    select-organism function, as follows:

    (select-organism :org-id "eco") to load EcoCyc, replace "eco" with "bsub" to load BsubCyc, etc.

    If you have a file-based database of a particular version, use select-organism as follows:

    (select-organism :org-id "bsub" :dbms-type :file :version "27.0")

 3. Run evaluate-predict-operons by pasting the following expression into the Lisp prompt:

    (evaluate-predict-operons :use-weights *no-expression-logistic-weights* :expression-data nil )

    which will run the operon predictor on the current organism, using appropriate regression weights
    and no-gene-expression data

    To run using E coli gene expression data

     (evaluate-predict-operons :use-weights *logistic-weights* 
                                 :expression-data (build-gene-expression-table-for-operons 
		                                   "SupFile3-ecoli-gene-expression.txt"))

  4. The accuracy, sensitivity and specificity scores will be reported at the end of the function output.


To replicate the model data for each model (with or without expression data) use
function generate-logistic-data by evaluating the following Lisp expression:

(generate-logistic-data :expression-values
			(get-rh-ecoli-expression-data "path-to-expression-data")
			:output-file "path-to-output-file")

The output file is a data matrix starting with column headers.  The
first column indicates whether a gene pair has experimental evidence
for or against being contained in an operon.  Pairs without
experimental evidence are not listed in this file.  The remaining
columns are states of the different features of the pair.  The column
headers are as follows:

      DISTANCE - Distance likelihood value based on binned distance between pair members 
      GEXPR - Correlation of gene expression values
      GO - GO biological process overlap
      CPX - Protein Complex -- are the adjacent genes are in the same protein complex?
      PWY - Shared pathway - both members of the gene pair in the same pathway
      TRN - Pathway/Transporter - one pair member is a transporter for substrates of a pathway the other member is assigned to
      UD-PWY - Up/Downstream shared pathway
      UD-TRN - Up/Downstream pathway/transporter
      LEN - If length(g1) >= .25 * length(g2)
      CHR4 - Gene symbol similarity
      NOMIX - Similar products (both protein, both RNA)
      GC-CNT - GC similarity 
      CODON - Codon usage using cosign similarity metric

The files used in the paper are SupFile1-pair-data-with-gene-expression.txt and
SupFile2-pair-data-no-gene-expression.txt.
