## Supplementary material for "Improved BioCyc Operon Prediction: Revisiting the Operon Prediction Problem": Performance of predictor for E coli and five other organisms

Supplemental Table 9

Performance of operon predictor across organism databases
  that provide experimental evidence for or against gene pairs being in an
  operon. The first line reports new predictor performance with gene
  expression data, the remaining lines use the new predictor
  without gene expression data. The second column
  reports counts of gene pairs that could be assigned to either be in
  an operon or not, based on experimental evidence, and the number of
  experimentally supported TUs (operons + single genes not occurring
  in an operon).
				                Accuracy		Sensitivity		        Specificity
Organism		exp pairs/TUs	New	Old	&New	Old	New	Old
E. coli K-12 MG1655(1)	1625/1428	83.099		83.490		82.544
E. coli K-12 MG1655(2)	1625/1428	82.547	79.55	82.863	76.622	82.101	83.78
Listeria monocytogenes 10403S	206/110	90.777	89.904	85.106	82.979	95.536	95.614
Streptomyces coelicolor A3(2)	188/110	80.319	83.511	82.105	81.053	78.495	& 86.022
Bacillus subtilis 168	182/88	79.670	80.769	77.982	77.064	82.192	86.301
Pseudomonas putida KT2440	65/26	70.769	70.769	82.051	79.487	53.846	57.692
Clostridioides difficile 630	65/17	83.077	75.385	 80.0	72.5	88.0	80.0
(1) Gene expression data used in model fit and prediction run
(2) No gene expression data used in either model fit or prediction run
