## Supplementary material for "Improved BioCyc Operon Prediction: Revisiting the Operon Prediction Problem": Summary of Experiments with Conditional Random Field models

This table compares performance of our CRF model with a L2-regularized logistic
regression model in 10-fold cross validation of segments. This is a much harsher
metric than binary accuracy because a model must exactly predict the entire operon
rather than pairwise in-operon relationships,  and it is generally harder to predict
longer operons.  We can see that CRF's appear better for predicting shorter operons
which make up the vast majority--transition probabilities may implicitly account for
length.  Concretely,  the BEGINING to END transition only appears in length-2 operons.
Overall, CRF appears superior by a margin of about 13.5% F1 measure.

		      R	     P		    F1
Len	 Count	 CRF	LRL2	CRF	LRL2	CRF	LRL2
 2	 190 	  0.679 	  0.579 	  0.717 	  0.458 	  0.697 	  0.512
 3	 81  	  0.543 	  0.346 	  0.512 	  0.326 	  0.527 	  0.335
 4	 61  	  0.508 	  0.410 	  0.554 	  0.510 	  0.530 	  0.455
 5	 30  	  0.500 	  0.567 	  0.750 	  0.607 	  0.600 	  0.586
 6	 19  	  0.684 	  0.526 	  0.619 	  0.667 	  0.650 	  0.588
 7	 10  	  0.400 	  0.200 	  0.267 	  0.250 	  0.320 	  0.222
 8	 5   	  0.400 	  0.200 	  0.250 	  0.167 	  0.308 	  0.182
 9	 6   	  0.333 	  0.333 	  0.333 	  0.667 	  0.333 	  0.444
 10	 1   	  1.000 	  1.000 	  0.500 	  1.000 	  0.667 	  1.000
 11	 2   	  0.000 	  0.000 	  0.000 	  0.000 	  0.000 	  0.000
 12	 2   	  0.000 	  0.000 	  0.000 	  0.000 	  0.000 	  0.000
 13	 1   	  1.000 	  1.000 	  0.333 	  0.500 	  0.500 	  0.667
 14	 0   	  0.000 	    N/A 	  0.000 	    N/A 	  0.000 	    N/A

Overall
	408	  0.593	  0.483	0.607	0.449	0.600	0.465
