## Supplementary material for "Improved BioCyc Operon Prediction: Revisiting the Operon Prediction Problem": List of NCBI SRR files used for gene expression analysis

### 1 Data for *Escherichia coli K12 M1655 Gene Expression Correlation*

Since McClure et al. (2013) did not report how datasets were assigned to experimental conditions, we list each SRA submission along with which of the 21 conditions (based on genotype, growth medium, time) we assigned it to. Conditions and replicates are as indicated in table 7 of supplementary material to McClure. Each NCBI Short Read Archive (SRA) submission was downloaded in June 2022. The SRA accessions were processed with Rockhopper 2.03 and the resulting gene expression data were used to calculate correlations, yielding our supplementary datafile correlations.txt.

| Experiment | Genotype / Conditions | SRA sequence ids |
|---|---|---|
| 1 | sgrS- control / 5 min after IPTG | SRR794821 |
| 2 | sgrS- + sgrS plasmid / 5 min | SRR794822 |
| 3 | sgrS- control / 10 min after IPTG | SRR794823 |
| 4 | sgrS- + sgrS plasmid / 10 min after IPTG | SRR794824 |
| 5 | sgrS- control / 20 min after IPTG | SRR794825 |
| 6 | sgrS- + sgrS plasmid/ 20 min after IPTG | SRR794826 |
| 7 | sgrS- control | SRR794827,SRR794828,SRR794829 |
| 8 | sgrS- + srgS+ plasmid | SRR794830,SRR794831,SRR794832 |
| 9 | sgrS- / minimal mops glycerol medium + $\alpha$MG | SRR794863,SRR794864,SRR794865 |
| 10 | sgrS- / minimal mops glycerol medium - $\alpha$MG | SRR794866,SRR794867,SRR794868 |
| 11 | WT / LB Medium + $\alpha$MG | SRR794833,SRR794834,SRR794835 |
| 12 | WT / LB Medium - $\alpha$MG | SRR794836,SRR794837,SRR794838 |
| 13 | WT / LB Medium + $\alpha$MG | SRR794839,SRR794840,SRR794841 |
| 14 | sgrR- in LB + $\alpha$MG | SRR794842,SRR794843,SRR794844 |
| 15 | WT / defined medium with glycerol + $\alpha$MG | SRR794845,SRR794846,SRR794847 |
| 16 | sgrS- / defined medium with glycerol + $\alpha$MG | SRR794851,SRR794852,SRR794853 |
| 17 | sgrS- / defined medium with glycerol - $\alpha$MG | SRR794854,SRR794855,SRR794856 |
| 18 | WT / minimal mops glycerol midium + $\alpha$MG | SRR794857,SRR794858,SRR794859 |
| 19 | WT / minimal mops glycerol midium - $\alpha$MG | SRR794860,SRR794861,SRR794862 |
| 20 | WT / minimal mops glycerol + 2DG | SRR794869,SRR794870,SRR794871 |
| 21 | WT / minimal mops glycerol midium - 2DG | SRR794872,SRR794873,SRR794874 |
