## Supplementary material for "Improved BioCyc Operon Prediction: Revisiting the Operon Prediction Problem": Script File for running Rockhopper for correlated expression analysis

#!/bin/sh
###  Instructions and utilities for working with Rockhopper
###    The inputs to Rockhopper are short-read data from multiple experiments,
###  and files describing the genome of that organism.
###    The output of Rockhopper is a file listing expression levels for each gene
###  across each of the experiments.  Our operon predictor uses this output file
###  to compute correlations in gene expression levels.

# How to run Rockhopper. (Rockhopper homepage: https://cs.wellesley.edu/~btjaden/Rockhopper/) 
#
# 1. Get a set of RNA-seq files for your organism, such as from NCBI SRA.  

# 2. Review the RNA-seq files and assign files to experiments.  If your set of files includes replicated conditions, 
#    Rockhopper does work with replicates.

# 3. Generate fna, ptt, and rnt files for your organism, or use the one on the Rockhopper site.
#    Put these files in a directory, one directory per organism.  These files describe the genome
#    sequence of the organisms, and its gene annotations.
#

#  We run Rockhopper with reference genomes (meaning the genome is already annotated),
#  so rockhopper is called the -g argument the directory containing the genome files (fna, ptt, rnt)
#  followed by a set of fastq files, with a set of experiments.

## Rockhopper requires Java (1.6 or better). and needs some things to be set up.
# Here is a sample script to run Rockhooper, with all
# 21 experimental conditions used to generate our expression dataset and E coli K-12 MG1655 genome files


export JAVA_HOME=/usr

if [ "${JAVA_HOME}" = "" ]; then
    echo 1>&2 Error: JAVA_HOME must be defined
    exit 127
fi

if [ "$WINDIR" = "" ]; then
    PATHSEP=':'
else
    PATHSEP=';'
fi


ROCKHOPPER_HOME="~/operon-prediction/data"  # put your datafiles in this directory
CLASS_FILES="~/operon-prediction-2021/Rockhopper-2.0.3/"  # points to the RockHopper class files
# PROGRAM_JAR="/homedir/midford/Projects/operon-prediction-2021/Rockhopper.jar"


#if ! [ -f ${PROGRAM_JAR} ]
#then
#    echo "Cannot find ${PROGRAM_JAR}"
#    exit 0
#fi

cd $ROCKHOPPER_HOME

# Sample script for running Rockhopper with Ecoli data as specified in the supplementary material from McClure et al. 2013

"${JAVA_HOME}/bin/java"  -Xmx2000m -cp $CLASS_FILES Rockhopper -g NC_000913 -v true srr794821.fastq srr794822.fastq srr794823.fastq srr794824.fastq srr794825.fastq srr794826.fastq srr794827.fastq,srr794828.fastq,srr794829.fastq srr794830.fastq,srr794831.fastq,srr794832.fastq srr794863.fastq,srr794864.fastq,srr794865.fastq srr794866.fastq,srr794867.fastq,srr794868.fastq srr794833.fastq,srr794834.fastq,srr794835.fastq srr794836.fastq,srr794837.fastq,srr794838.fastq srr794839.fastq,srr794840.fastq,srr794841.fastq, srr794842.fastq,srr794843.fastq,srr794844.fastq srr794845.fastq,srr794846.fastq,srr794847.fastq srr794851.fastq,srr794852.fastq,srr794853.fastq srr794854.fastq,srr794855.fastq,srr794856.fastq srr794857.fastq,srr794858.fastq,srr794859.fastq srr794860.fastq,srr794861.fastq,srr794862.fastq srr794869.fastq,srr794870.fastq,srr794871.fastq srr794872.fastq,srr794873.fastq,srr794874.fastq
